# High throughput mutational scanning of a protein via alchemistry on a high-performance computing resource

**DOI:** 10.1101/2024.08.20.608765

**Authors:** Tandac F. Guclu, Busra Tayhan, Ebru Cetin, Ali Rana Atilgan, Canan Atilgan

## Abstract

Antibiotic resistance presents a significant challenge to public health, as bacteria can develop resistance to antibiotics through random mutations during their life cycles, making the drugs ineffective. Understanding how these mutations contribute to drug resistance at the molecular level is crucial for designing new treatment approaches. Recent advancements in molecular biology tools have made it possible to conduct comprehensive analyses of protein mutations. Computational methods for assessing molecular fitness, such as binding energies, are not as precise as experimental techniques like deep mutational scanning. Although full atomistic alchemical free energy calculations offer the necessary precision, they are seldom used to assess high throughput data as they require significantly more computational resources. We generated a computational library using deep mutational scanning for dihydrofolate reductase (DHFR), a protein commonly studied in antibiotic resistance research. Due to resource limitations, we analyzed 33 out of 159 positions, identifying 16 single amino acid replacements. Calculations were conducted for DHFR in its drug-free state and in the presence of two different inhibitors. We demonstrate the feasibility of such calculations, made possible due to the enhancements in computational resources and their optimized use.

## I. INTRODUCTION

Bacteria constantly evolve to adapt to their surroundings through genetic mutations. These mutations, occurring at the gene level, can affect proteins that are essential for the organism’s survival. With natural selection, these mutations, even if they arise in extreme environments, can become permanent. This process lies at the core of the antibiotic resistance crisis which is among the most significant public health challenges today.

In our project that we conducted on the Karolina cluster, we concentrated on the enzyme DHFR, responsible for converting dihydrofolate (DHF) to tetrahydrofolate (THF) (Figure 1). THF is critical for DNA, RNA, and amino acid synthesis, making DHFR ubiquitous in all organisms. The resistance of *E. coli* DHFR to the drug trimethoprim (TMP), which competes with DHF for binding at the same site, has been extensively studied through systematic experiments, revealing a series of enduring mutations [1]. Some mutations may be temporary, appearing at certain stages but not in the final colonies while others are fixed early on [2]. On the experimental side, the advancements in molecular biology tools over the past decade have enabled deep mutational scanning (DMS) of proteins [3]. DMS libraries of *E. coli* DHFR now exist, with fitness quantified under different conditions and mapped to specific protein locations [4]. However, quantifying fitness poses challenges, as contributions can arise from folding pathways in addition to those directly related to protein function.

**Fig. 1.**
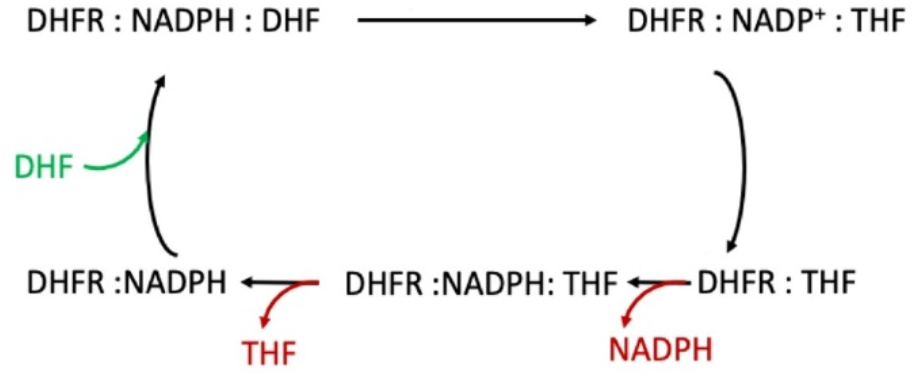
Five-step kinetic cycle of DHFR.

In our previous research, we have conducted molecular dynamics (MD) simulations of appropriate duration for specific mutants in both their DHF and TMP-bound states. Initially, we focused on understanding the molecular-scale mechanism of the frequently occurring L28R mutant [5]. This allowed us to elucidate the essential characteristics of the actions taken by various mutants, all of which were confirmed through experimental validation [2]. Utilizing the insights gained from the mechanism of the L28R mutant, we devised a derivative of TMP (4’-deoxy trimethoprim) that diverts evolutionary trajectories towards less harmful outcomes [6]. More recently, we have employed hydrogen bond dynamics and side-chain relaxations on nanosecond time scales to automatically uncover the atomistic-level mechanisms involved in the emergence of resistance [7].

In most cases, molecules with competitive binding are distinguished by changes in free energy, which in turn affect binding probabilities. However, since the mutants we investigated remain functional, the free energy differences due to these single mutants are usually small. Additionally, solvent effects contribute significantly to entropy/enthalpy compensation [8]. Consequently, coarse-grained free energy methods employing implicit solvation models are inadequate for computationally describing the observed effects.

In this study, we assess the fitness at the molecular level for all single mutants of DHFR at 33 positions. For this purpose, we utilized the free energy perturbation (FEP) approach, which currently yields values within 1 kcal/mol of experimentally measured binding free energies [9]. We constructed a library of mutants for three systems; DHFR in its drug-free from (apo), TMP-bound, and 4’DTMP-bound forms. By using suitable thermodynamic cycles, we predicted the change in the free energy cost of the mutations. To achieve this task, we constructed a pipeline for generating the systems of interest and analyzing the large amount of output.

The information collected in this study has the potential to contribute to the scientific literature in fundamental ways: (1) Experimental findings regarding DHFR’s drug resistance may be evaluated through the structural changes of the enzyme; (2) contribution of protein thermodynamics to drug resistance may be systematically studied; (3) findings may guide future work in developing new and potent TMP derivatives.

## II. METHODS

### A. Thermodynamic cycles

For understanding the relative binding between the drug TMP and the inhibitor 4’DTMP, two thermodynamic schemes are constructed (Figure 2).

**Fig. 2.**
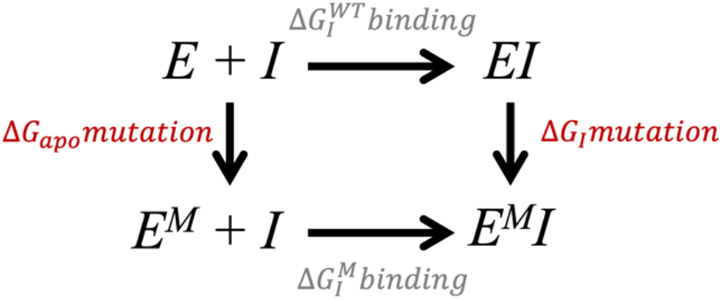
Horizontal lines represent binding events (gray), while vertical lines depict mutations on the enzyme (red).

To calculate the relative binding energy of 4’DTMP over TMP, we first need to understand the relation between mutational energies and the binding energies. *M* denotes the presence of a mutation in the protein, and *I* depicts the inhibitor (drug). Thus, the thermodynamic scheme of either TMP or 4’DTMP can be written as:

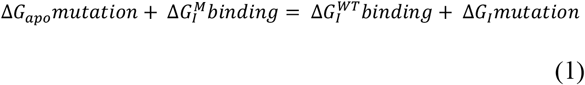

When we collect the mutational energies on one side the equality becomes:

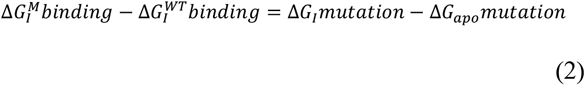

For calculating the relative binding free energies, we write equation (2) both for 4’DTMP and TMP. Then, by subtracting and rearranging the terms, we obtain:

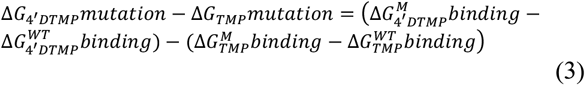

The last term is an additive constant which is the same for all systems and is ignored in the interpretation of the results. This set-up provides a means to compare the effectiveness of the two drugs in the presence of a mutation.

### B. System set-up

To calculate the free energy differences across the apo, TMP-bound, and 4’DTMP-bound systems, we first conducted 1 μs-long molecular dynamics (MD) simulations for each condition, from which the structures were obtained. MD simulations were performed by using NAMD [10] program. CHARMM36 force-field [11] was used for topologies and parameters. The dimensions of the water box were set to 62 × 68 × 60 Å, including a minimum of 10 Å TIP3P water layer in all directions. K^+^ and Cl^-^ ions were added to neutralize charges and maintain a 150 mM physiological ionic strength. NADPH was included in all simulations. The systems were minimized for 10 000 steps. The Particle Mesh Ewald method was employed for calculating long-range electrostatic interactions with a cutoff of 12 Å and a switching distance of 10 Å. The RATTLE algorithm was applied, and the Verlet algorithm was used with a time step of 2 fs. Temperature control at 310 K was achieved through Langevin dynamics, featuring a damping coefficient of 5 ps. Pressure was maintained at 1 atm, regulated by the Langevin piston. The production equilibrium simulations were performed for 1μs, and trajectories were stored every 10 ps.

### C. Selection of initial conformers for FEP calculations

We utilized different metrics to obtain structures from MD trajectories, such as solvent accessible area (SASA), hydrogen bonds between protein-ligand and evolution of conformational cluster populations during simulations (Figure 3). The number of hydrogen bonds formed between a ligand and a protein can provide valuable insights into the stability and dynamics of the drug-protein interaction. Therefore, structures with a higher number of hydrogen bonds were considered more stable and suitable for further analysis. SASA also helps discriminate the compactness and the stability of the structure. By analyzing the changes in SASA over time, valuable insights can be gained into the protein’s structural dynamics, including conformational changes. Therefore, by using this metric, it is possible to distinguish structures across different conformational states. The evolution of conformational cluster populations over time during the simulation was observed by using CPPTRAJ [12]. These clusters, representing molecular structures with similar shapes or conformations, offer insights into the dynamic behavior of proteins, revealing their structural changes throughout the simulation duration. Based on these analyses, three conformations (utilized as replicas) were selected from each condition for FEP simulations. Due to the restricted amount of computation time available to us on the Karolina cluster, we were able to complete 33 positions of the total of 159 positions with 16 replacements; PRO, GLY and another selected for each position were excluded. During the selections, the most chemically similar variant of the WT strain at that position was discarded.

**Fig. 3.**
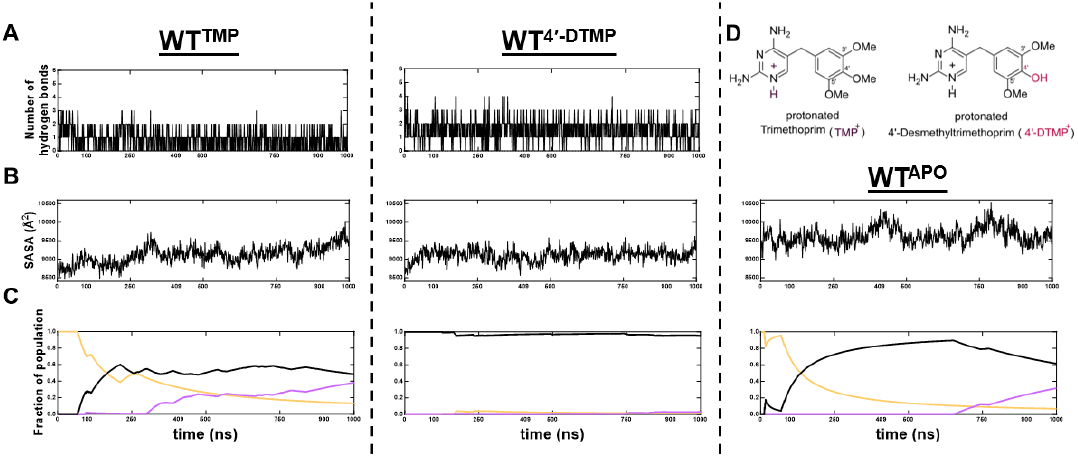
Different parameters used to select conformations for FEP simulations. (A) Number of hydrogen bonds between inhibitor and DHFR along the trajectory; (B) SASA of apo, TMP-bound, 4’DTMP-bound systems during simulations; (C) development of conformational clusters with time calculated via CPPTRAJ; (D) chemical structures of the two drugs.

### D. Alchemical free energy perturbation calculations

Alchemical FEP calculations involve the application of Zwanzig’s method [13]. All calculations employ the CHARMM36 force field [11] and are made using NAMD3 [14] program. With the development of NAMD3, it has become possible to run FEP simulations using a GPU and a lower number of CPU cores (8 – 16), which allows us to plan for running a large number of FEP simulations [15]. The protein is again soaked in a solvent box such that there is at least a 10 Å layer of solvent in every direction from any atom of the protein to the edge of the box. K^+^ and Cl^-^ ions are added to neutralize charges and maintain a 150 mM physiological ionic strength. The protocol explained in detail in the literature was used [16]. Briefly, particle mesh Ewald sum was employed to control the electrostatics. Langevin piston was used for pressure control at 1 atm, and the temperature was maintained at 310 K as in the main MD simulation run. Switching was implemented with a cutoff of 13 Å and a switching distance of 10 Å. The timestep was set to 2 fs. The SHAKE algorithm was used for all bonds. Systems were minimized for 500 steps. The systems were run for *λ* = 32 windows in both forward and backward directions. To address singularities encountered during Lennard-Jones potential calculations, a soft-core potential was incorporated to truncate the function for *λ* = 16 – 32. Sampling was performed from the classical MD simulations as described in the previous subsection to enhance coverage of potentially available conformational states. MD simulations were carried out in each window for 0.2 ns, with the initial 0.1 ns of each window discarded for equilibration. Results were analyzed using Bennett-acceptance ratio (BAR) method [17]. Three replicate FEP simulations for each mutational change are concatenated to reduce the error.

### E. Charge corrections

In FEP simulations, during the perturbation process, the wild-type amino acid gradually disappears from its sequence position while the mutated amino acid simultaneously emerges in its place. The charge characteristics of both the wild-type (WT) and mutated amino acids are crucial during this transformation. As the system’s charge shifts due to the mutation – such as transitioning from negative to positive or neutral to positive – K+ and Cl− ions are removed to maintain the system’s overall neutral charge. For example, when changing from a positively charged amino acid to a negatively charged one, two K+ ions are removed. Conversely, when shifting from a negatively charged position to a positively charged mutation, two Cl− ions are removed to neutralize the system. In cases where transitions occur between neutral and charged states, only one ion is removed. The ions selected for removal are chosen randomly, ensuring they are located more than 20 Å from the protein.

### F. Positions selected

Due to constraints in computational resources allocated to our project on the Karolina cluster, we adopted a selective approach in determining the positions to be investigated. These selections were informed by prior studies on DHFR. Initially, attention was directed towards positions identified through morbidostat experiments under trimethoprim selection, aiming to elucidate their potential impact on 4’DTMP binding (positions 5, 20, 26, 28, 30, 94, 98, and 153) [2]. Subsequently, emphasis was placed on residues situated within cryptic sites identified in our earlier work (positions 59, 69, 70, and 71) [7]. Furthermore, positions delineated as hotspots in the literature [18] were included in the analysis (positions 16, 33, 36, 48, 50, 73, 80, 83, 114, 121, 124, 127, and 131-132). Lastly, hinge regions (positions 87-88, 106-107) of DHFR were incorporated into the study [19], along with mutations (positions 16, 19 [20], and 42 [21-24]) known to impact catalysis. Additionally, each position was mutated to 16 of the 19 possible positions, meaning that three replacements were omitted in each case. This omission was due to our optimization of cluster usage to multiples of 8, which allowed us to make the best use of the computing time available to us. These included mutations involving proline residues which are not reliably treated with FEP simulations at this time due to the unique situation of the ring that is bonded to the backbone being created or annihilated. In all cases, the glycine residue was also omitted due to its small size, which can lead to large error bars in the amino acid replacement process of FEP calculations[25]. The third position to be omitted was chosen based on the chemical properties of the amino acid being replaced. Hence, a total of 33 amino acid positions were selected for single mutational scanning using FEP simulations.

### G. Computer time availability

In this project, we were allocated 3,000 node hours on the Karolina-GPU high-performance computing (HPC) facility. The GPU-accelerated nodes of the Karolina HPC consist of 2 × AMD EPYC™ 7763, 64-core, 2.45 GHz processors (a total of 128 CPU cores) and 8 × NVIDIA A100 GPUs, providing an optimal environment for conducting large-scale FEP simulations. We utilized the NAMD3 software to conduct FEP simulations, encompassing three stages: minimization, forward, and backward. These subprocesses were executed sequentially, with groups of eight FEP simulations running in parallel. Each FEP simulation required the use of 16 CPU cores and one GPU from the Karolina-GPU HPC facility. This approach enabled the completion of 48 FEP simulations per day, thereby facilitating the investigation of mutational perturbations at three amino acid positions in one day.

## II. RESULTS And DISCUSSION

### A. Pipeline developed for multiple free energy perturbation simulation for single-mutational scanning of a protein

We used NAMD3 software to conduct 1,584 simulations (33 positions × 16 mutations × 3 replicates) for each of the apo, TMP-bound, and 4’-DTMP-bound forms, totaling 4,752 FEP simulations. Each FEP simulation consisted of three subprocesses: minimization, forward, and backward. We executed these three subprocesses sequentially, while concurrently running groups of eight FEP simulations in parallel. Tcl scripts were utilized to prepare each simulation on local computers. To execute this complex process on HPC, the submission file for workload manager was generated via a homemade Python script. To perform a single mutational scanning with FEP of a position for apo and ligand-bound forms, we uploaded ∼1,500 files (1 GB) to the HPC. After the simulation execution, ∼7,700 files (36 GB) were downloaded. File transfers and consistency checks were performed by assessing log files via terminal commands. We revised our job submission script due to a change in the workload manager, causing a delay in the completion of our simulations. Possible errors during the simulation were investigated through daily spot checks of simulation files.

Each FEP simulation utilized 16 CPU cores and 1 GPU from the Karolina-GPU HPC, enabling the investigation of mutational perturbations at one amino acid position per day. For each FEP simulation, we produced an equivalent of 125 ± 22 ns-long simulations per day (ns/day) in the Karolina-GPU queue. The efficiency of the simulations varies based on the initial structures and mutations. Each simulation for a positional change is approximately 20 ns long, and it takes 0.16 days to complete one FEP simulation. By running eight FEP simulations in parallel across multiple nodes, and accounting for the time lost between each batch of simulations, we were able to perform 144 (1 position × 16 mutations × 3 replicates × 3 forms of DHFR) simulations per day.

The pipeline is designed for general usage to perform mutational scanning of protein structures using FEP simulations. The scripts are easy to modify, and the generation of submission files is well documented on our GitHub page. By using the provided codes, it is possible to perform a large number of FEP simulations on a variety of HPC systems with different configurations.

We note that while this protocol uses 6.4 ns for each alchemical transformation (ca. 20 ns per position in three replicates), one can modify the length of the simulations in proteins where the mutational effects are known to induce significant conformational changes in the protein, due to, e.g., a loop changing orientation preference or allosteric effects. This preference will depend on the availability of the computational resources.

### B. Information on benchmarks and energies for FEP simulations conducted based on amino acid charges

We grouped our FEP simulations based on the charge differences between wild-type and mutated amino acids. For mutations involving a two-charge difference (e.g., negative to positive, +2, or positive to negative, -2), two ions are removed. For mutations involving a one-charge difference (e.g., neutral to positive, +1, neutral to negative, -1, and their opposites), only one ion is removed. No ions are changed for transformations involving residues with the same charge type.

To investigate the effect of an increased number of changes in the system on computing time, we analyzed the benchmark results of NAMD3 software and found that, on average, our FEP simulations run at approximately 120 ns per day (Figure 4). The number of atoms and interactions (van der Waals and Coulomb) affect the benchmark results; thus, the changing nature of the amino acids and the removal of different numbers of ions influence computation time. Mutations applied to neutral and positive residues exhibit similar behavior in terms of computational time. However, negative wild-type amino acids show greater deviation in benchmark values. Although both the charge and position of the residue influence computational time, a negative wild-type amino acid may have fewer interactions compared to other mutations, leading to the lower tail in the distribution of computational time usage.

**Fig. 4.**
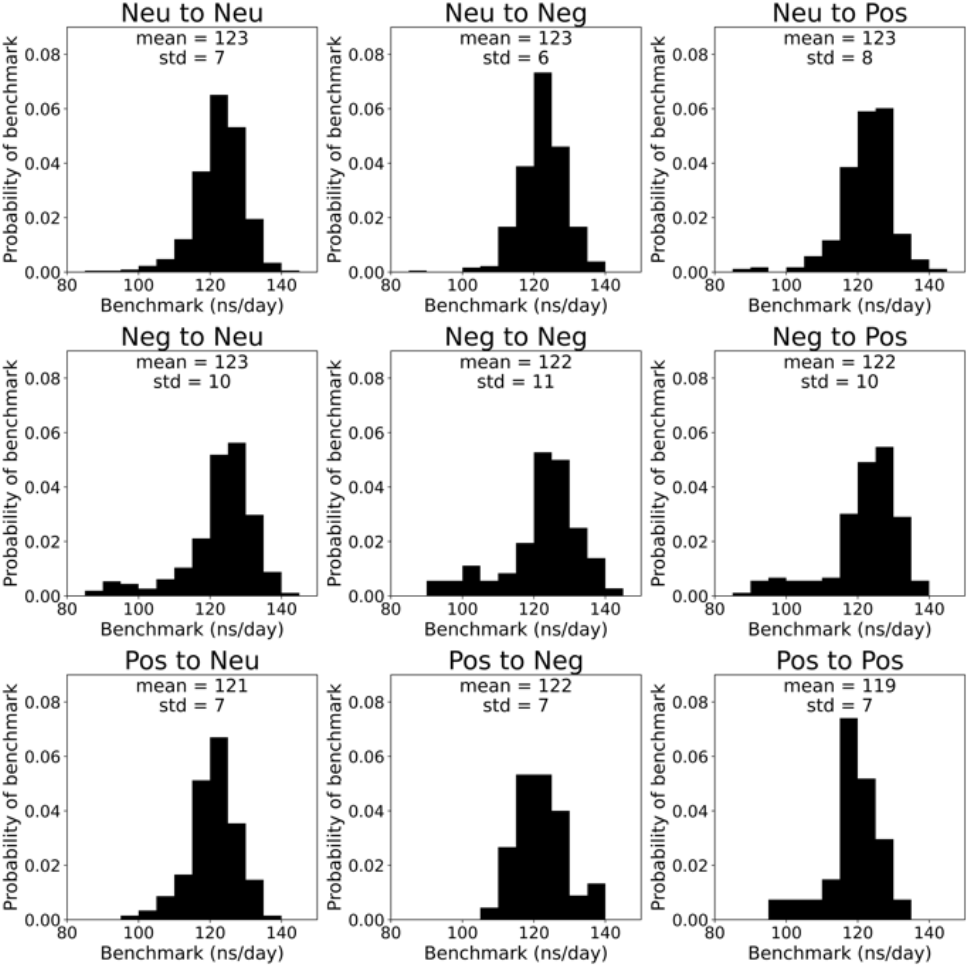
The probability of the benchmark (ns/day) was calculated for each forward process of the FEP simulations. The benchmark values calculated for each simulation by the NAMD3 software were utilized across all 4,752 simulations. The simulations were clustered based on the charge of the wild-type and mutated amino acids.

We further investigated the mutation energies for the apo and TMP/4’DTMP-bound forms. Our results indicate that mutations of the same type cause only minuscule changes on average, with a slight increase observed in neutral to negative mutations (Figure 5). The highest energies are associated with mutations from negative/positive to neutral and from negative to positive. Interestingly, positive to negative mutations exhibit only half the energy changes compared to negative to positive mutations, suggesting that the position of the residues also significantly impacts the energy changes (Figure 5). Additionally, favorable mutations with negative energy values occur only when mutating neutral amino acids (Figure 5).

**Fig. 5.**
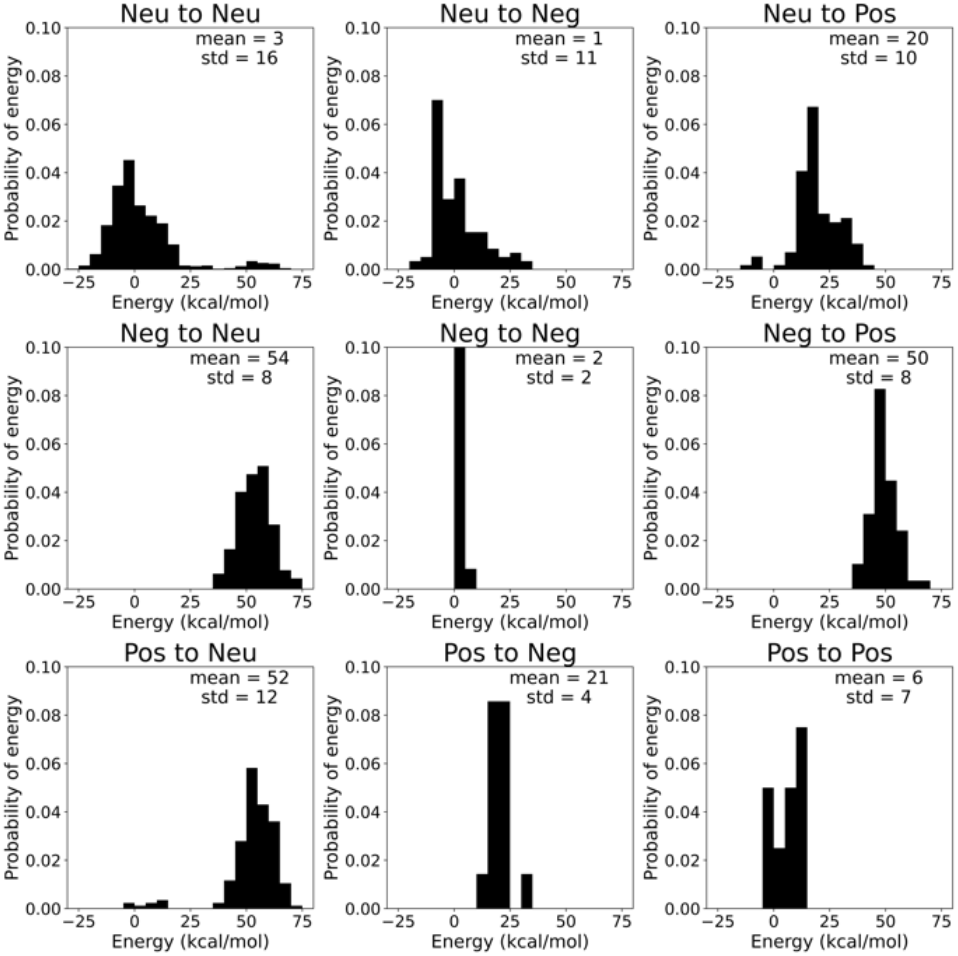
The probability distribution of the mutation energy (kcal/mol) was calculated for each mutational change. The energy values calculated for each mutational change, calculated using the BAR method, were utilized for 1,584 mutation energies. The energy values were clustered based on the charge of the wild-type and mutated amino acids.

The accuracy of a free energy calculation can be assessed by the reported variance in the reported free energy change value, with the expected variance for computing mutation or relative binding energies being approximately 1 kcal/mol [9].

To evaluate the accuracy of the BAR-analyzed mutational energies for both apo and TMP/4’DTMP-bound forms, we visualized the distribution of variances (see Figure 6). We found that the average variance for all mutations is significantly lower than 1 kcal/mol. The highest variances are observed for charge-changing mutations (positive to negative and negative to positive), as these involve altering both the amino acid and two ions, leading to a higher accumulation of error in the FEP simulations (see Figure 6). Note that these variances are calculated separately for the mutational changes of the apo and TMP/4’DTMP-bound forms. When calculating relative binding energies and summing these values, the variance increases, though it remains around 1 kcal/mol.

**Fig. 6.**
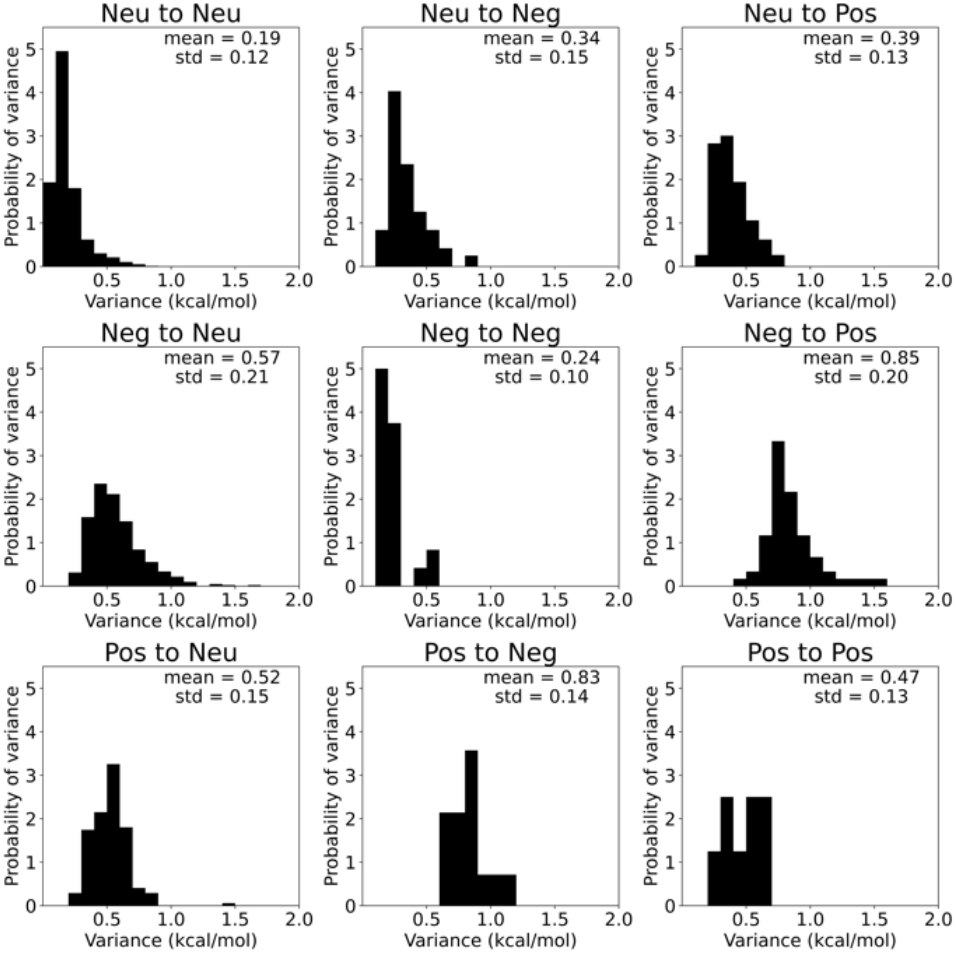
The probability distribution of the variance in the mutational energy cost (kcal/mol) was calculated for each change. The variances calculated for each mutational change by BAR method were utilized for 1,584 mutation energies. The variance values were clustered based on the charge of the wild-type and mutated amino acids.

### C. Relative binding energies explains the mutational landscape of binding

To further understand the effective drug, we calculated the relative binding energies. The ΔΔ*G* values calculated according to equation 3 are displayed in Figure 7. Negative energy values indicate favorable binding to 4’DTMP, while positive energy values correspond to favorable binding of TMP. When comparing the relative binding free energies of all positions for sixteen mutants, we have found that 4’DTMP is more effective at twenty-six out of the thirty-three positions typically surrounding the active site and dissipating throughout the enzyme. The positions that display mixed behavior (6 out of 33) may belong to a pathway through the signaling of I5 to cofactor site R98 connected with the constituents of FG loop. Finally, the only position where TMP is more effective is D69, an important player on the formation of the cryptic site we discovered in our previous work [7].

**Fig. 7.**
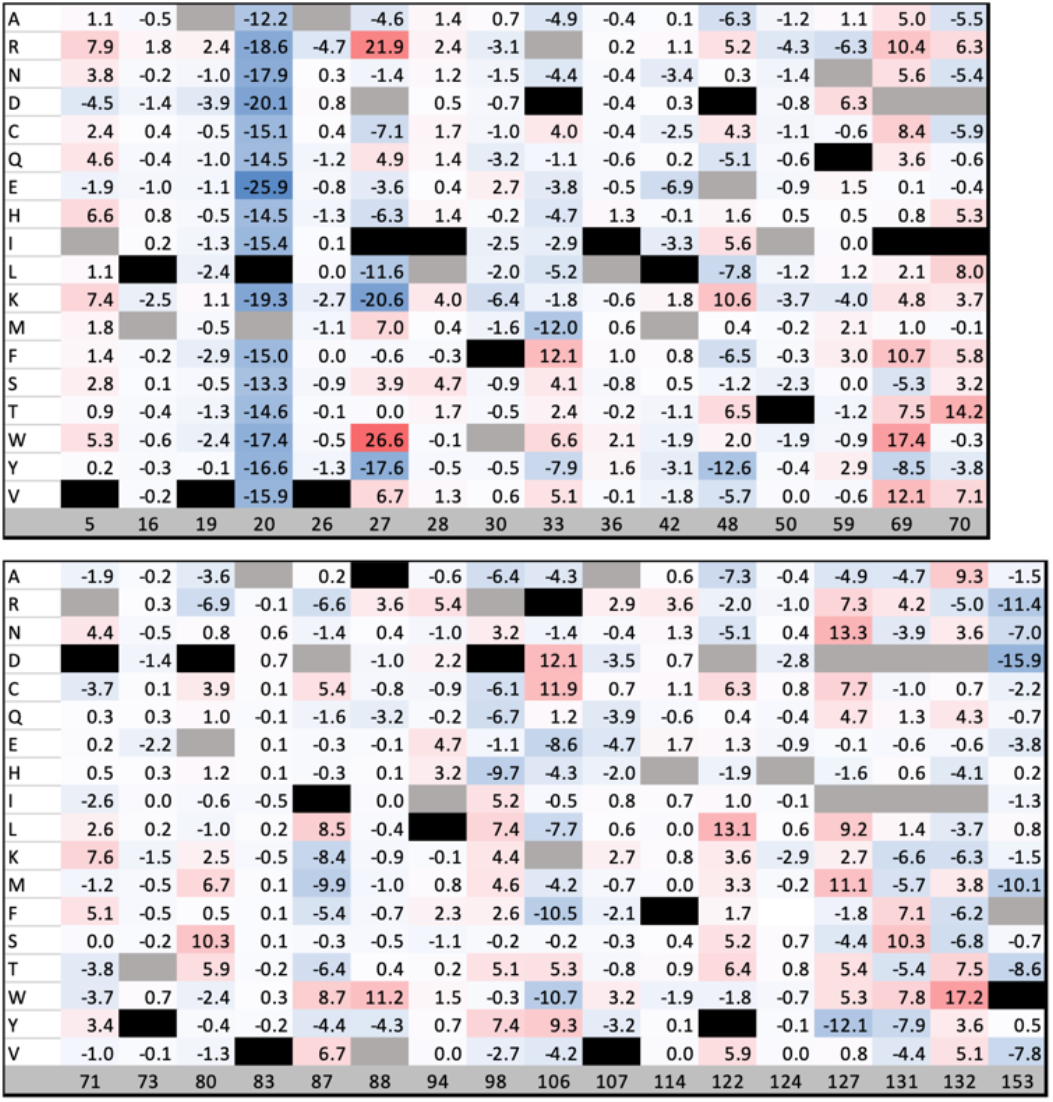
Color coded representation of calculated free energy differences for 4’DTMP and TMP (Equation 4). Blue indicates the mutants that have higher affinity toward 4’DTMP, and red towards TMP. Grey indicates the position of the amino acids in the WT protein, while black mutations are those that are omitted in this study.

Our general observations are as follows: (i) Mutations in the binding site that provide access to the trimethoxy tail exhibit a greater affinity towards 4’DTMP; otherwise, TMP is stabilized. The catalytic loop residue M20 is special in that its mutation always favors 4’DTMP. (ii) Mutations at our previously identified cryptic site are sensitive, especially at positions 69 and 70 which mostly increase preference to TMP binding. (iii) Allosteric communication with distal sites permeates throughout DHFR, manifesting alternative communication pathways in the catalyst. Some positions that are strongly involved in this type of mechanism are D87, K106, D122, D131, D132 which were also delineated as hotspots in the literature [18] and F153 which was found to arise frequently in morbidostat experiments [2]. Finally, we note that FEP used for relative binding energy differences assumes that the changes do not affect the binding mode of the ligand. For the drugs used in this work, TMP and 4’DTMP, which have high (nanomolar) binding affinity, this is not a problem but should be further explored on a case-by-case basis, especially if the experimental predictions and the computed differences diverge.

We also computed the variances of relative binding energies for all single mutational changes across 33 positions, and our results indicate that the error values are, on average, lower than 1 kcal/mol (Figure 8). Overall, our FEP simulations investigating the relative binding energy changes upon single mutations demonstrate the desired precision [9].

**Fig. 8.**
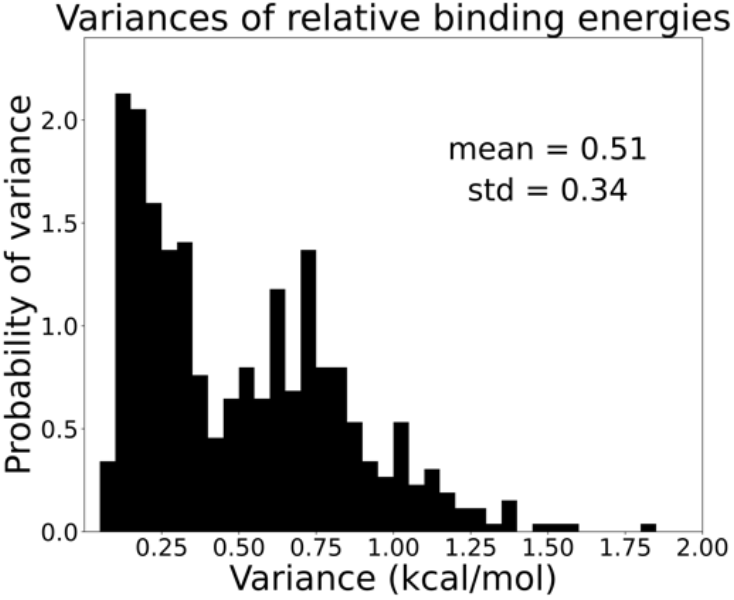
The probability distribution of the variance of relative binding energy calculated for all single mutations across 33 positions.

We further investigated the minimum and maximum relative binding energies at each position to identify the most effective mutant with favorable binding. On average, the minima and maxima ranged from -4 to 4 kcal/mol across 33 positions. We visualized the positions with the highest energies by selecting those with absolute values greater than 4 kcal/mol (Figure 9). We found that 25 out of 33 positions exhibited high relative binding energies. Notably, position M20 favors only 4’DTMP, while other positions display both negative and positive relative binding energies, indicating that specific mutations can have significantly different effects depending on the position.

**Fig. 9.**
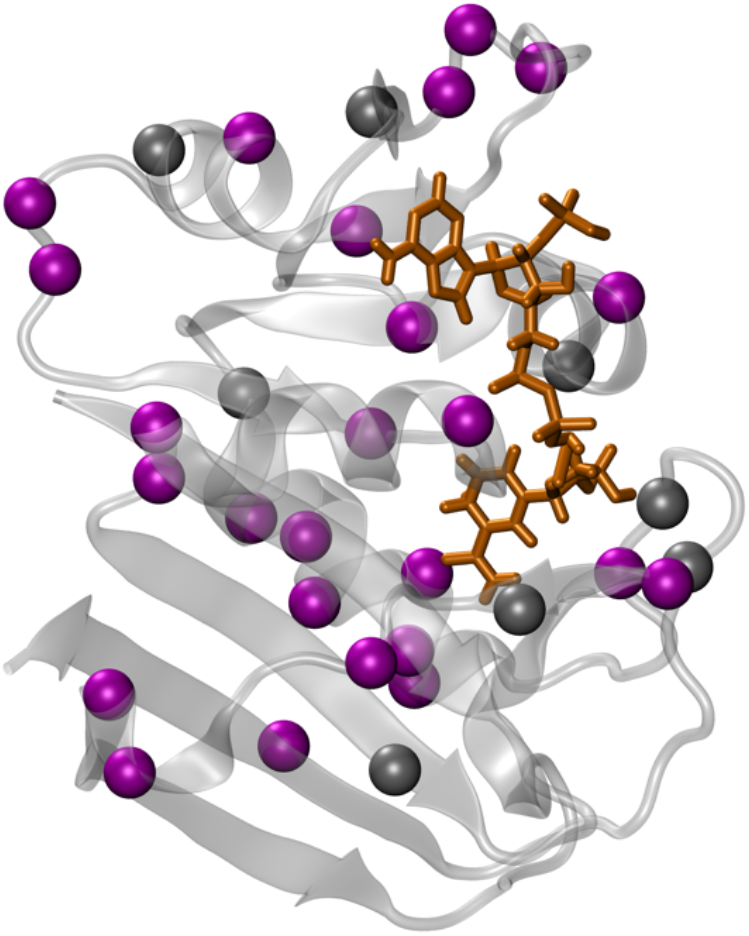
The residues with the highest relative binding energies (25 out of 33 positions) are visualized on the apo DHFR structure. Residues with a relative binding energy greater than an absolute value of 4 kcal/mol are colored magenta, while other amino acids are shown in gray. NADPH is shown in orange.

This work lays the foundations for assessing the effects of point mutations on protein-drug affinities using alchemistry on HPC resources. With the increase in the available experimental data in this realm, it is imperative that automated schemes such as the one proposed here to set up MD simulation systems and codes for analyzing the large amount of collected output be available to researchers in a wide range of fields.

## Data and Code Accessibility

The scripts are shared on our group’s GitHub page: https://github.com/midstlab/FEP_mutational_scanning

